# The head-direction signal is generated by multiple attractor-like networks

**DOI:** 10.1101/2025.09.10.675474

**Authors:** Guillaume Viejo, Sofia Skromne Carrasco, Adrien Peyrache

## Abstract

The generation of the head-direction (HD) signal, a cornerstone of the brain’s navigation system, is classically attributed to the lateral mammillary nucleus (LMN) and associated brainstem structures, where attractor-like dynamics are thought to maintain population coherence. Here, using multi-site recordings and optogenetic perturbations along the mammillary–thalamic–cortical pathway, we demonstrate that HD neurons in the anterodorsal nucleus of the thalamus (ADN) maintain coherent population dynamics even when their LMN inputs become decorrelated during non-REM sleep. These findings reveal that thalamic coherence does not strictly depend on structured input from the LMN; instead, it can emerge from local thalamic processes involving shared inhibition and non-linear responses. Together, our findings reveal a previously unrecognized, state-dependent shift in the circuit organizing the HD signal, establishing the thalamus as an active substrate capable of independently sustaining internal representations across brain states.

## Introduction

Continuous attractor networks are believed to support various cognitive functions, from working memory to spatial representations, yet the neuronal dynamics and circuits supporting these dynamics in vivo remain unclear ^1–4^. One example of such networks is the head-direction (HD) system ^5–7^, a crucial signal for navigation ^8–11^. It is represented by HD cells, which each fire for a specific direction of the animal’s head, and is transmitted to the cortex by the anterodorsal nucleus (ADN) of the thalamus where a vast majority of neurons are modulated by HD ^12,13^. ADN-HD cells maintain their mutual coordination during sleep ^2,13,14^, when sensory inputs are virtually absent, both during rapid eye movement (REM) sleep (an activated state like wakefulness) and non-REM sleep (a deactivated state). This sustained coherent activity across brain states is consistent with an attractor-driven system ^2^.

This coherent nature of ADN activity raises a fundamental question: where is the HD attractor-like activity generated? Prevailing models identify the lateral mammillary nucleus (LMN), the primary input to the ADN, as the central generator of the HD signal ^5,15,16^. In this view, the thalamus acts as a relay, passing the LMN’s attractor dynamics to the postsubiculum (PSB) and broader cortical networks ^7,13,17^. The PSB, in turn, provides feedback to the LMN ^18,19^.

Here, we investigated the ensemble organization of LMN activity across brain states and its relationship to ADN dynamics. We show that during non-REM sleep, coordination among LMN-HD cells is significantly degraded, yet simultaneously recorded ADN neurons remain as coherent as during wakefulness. Furthermore, while silencing cortical feedback (PSB) abolished the remaining LMN coordination, ADN dynamics remained intact. ADN neurons showed strong non-linear responses, a property that, together with putative shared inhibition, is sufficient to maintain coherent activity in a noise-driven regime. Our findings establish that the HD signal is supported by multiple attractor-like stages, revealing the thalamus as an independent and active node.

## Results

### Coherence of HD cell population activity is maintained in the ADN during non-REM, but not in the LMN

To investigate the origin of structured activity in the ADn during sleep ^2,13,14^, mice were chronically implanted with high-density silicon probes targeting the LMN (*n* = 11), ADn (*n* = 4), or both (*n* = 8) (Figure 1A–D; Supp. Figure 1). We recorded a total of 546 HD cells in the LMN and 265 in the ADn (Supp. Figure 2A-B). Mice explored a circular environment and were allowed to sleep before and after exploration.

**Figure 1.**
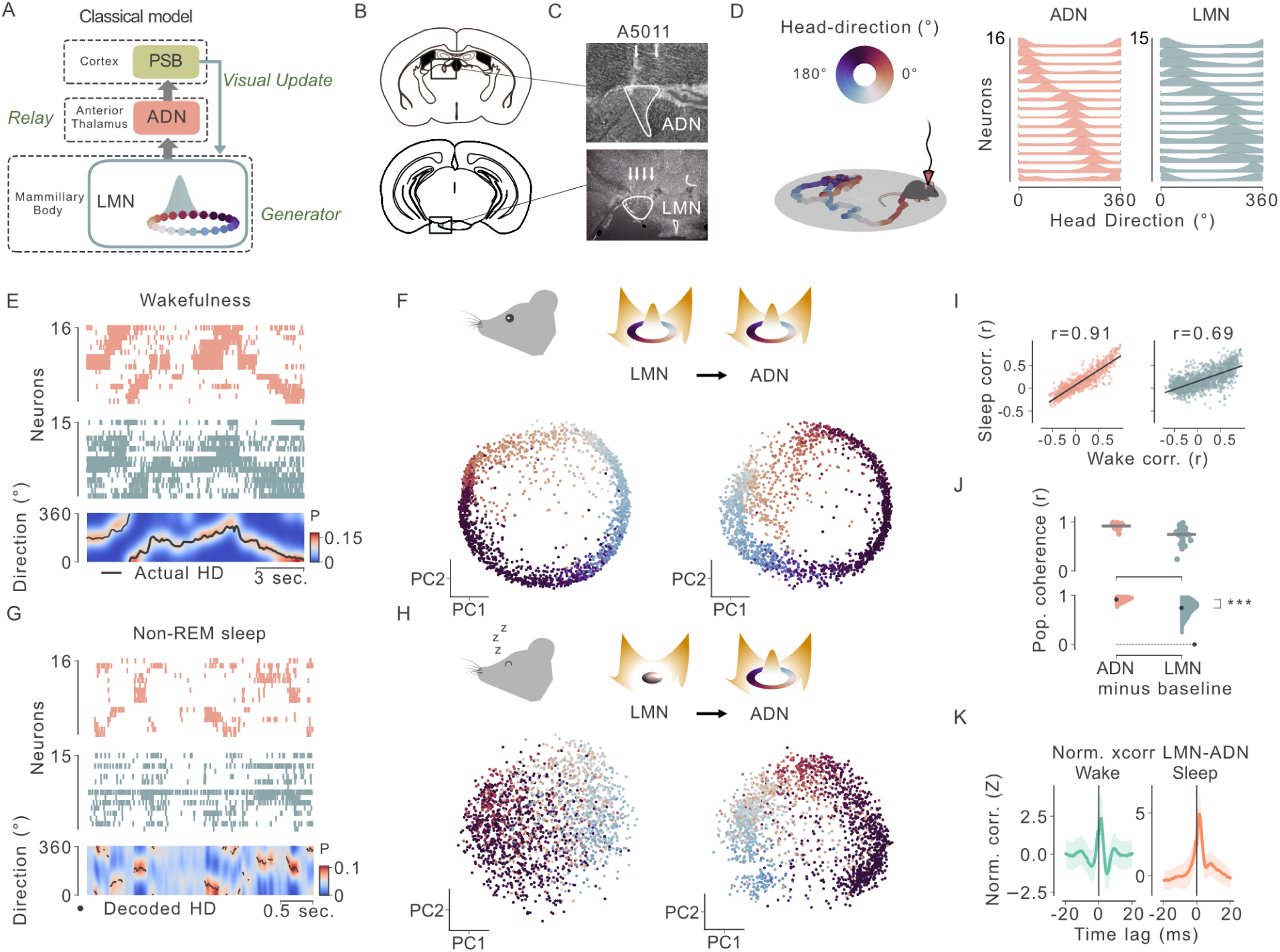
Decreased HD cell population coherence in LMN, but not ADN, during non-REM. **(A)** Schematic of the classical model for HD signal generation involving the cortex (postsubiculum, PSB), anterodorsal nucleus of the thalamus (ADN), and lateral mammillary nucleus (LMN). **(B)** Coronal diagrams showing LMN (bottom) and ADN (top) location. **(C)** Histological verification of probe locations (arrows) in the ADN (top) and LMN (bottom). **(D)** Left: Example trajectory of a freely moving mouse in a circular environment. Circular coloring indicates head-direction at each position. Right: Tuning curves of HD neurons from ADN (left) and LMN (right) sorted by preferred direction for one example session. **(E)** Top: Raster plots of HD cell spiking activity in ADN (orange) and LMN (blue) during wakefulness. Bottom: Decoded HD signal, showing clear activity packets during wakefulness in both structures. Black line indicates actual head-direction. **(F)** Projections of neural population activity during wakefulness for LMN (left) and ADN (right). Points are color coded by head-direction. **(G)** Same as E for non-REM sleep. Bottom: Note the clear activity packet in ADN, unlike LMN. Black dots show the decoded HD signal (i.e. angle of maximum decoding probability). **(H)** Same as F during non-REM sleep. Note the lack of ring structure for LMN activity compared to ADN. **(I)** Correlation of pairwise coupling between wake and sleep for all HD cell pairs in ADN (left) and LMN (right). **(J)** Population coherence during non-REM, quantified as pairwise coupling minus a spike-shuffled baseline, was significantly higher in ADN compared to LMN (Mann–Whitney; U=111; p<0.001; n_ADN_=24; n_LMN_=35 sessions). **(K)** Normalized cross-correlograms of LMN→ADN cell pairs during wakefulness (green) and non-REM sleep (orange) show strong short-latency peaks consistent with monosynaptic influence in both states.

As expected, during wakefulness, HD cell population activity accurately tracked the animal’s head direction, forming a well-defined “activity packet” (Figure 1E). Low dimensional projections of the neural activity (see Methods) during wakefulness revealed a ring structure typical of continuous attractor dynamics for both LMN and ADN (Figure 1F). The structure of HD cell activity was maintained in both structures during REM sleep (Supp. Figure 2C), suggesting that during activated states of the brain, similar processes are at play in the generation of a coherent HD signal ^7,13,14^.

During non-REM sleep, however, dynamics differed strikingly between the two structures (Figure 1G). As previously reported ^2,13,14^, the ADN-HD cell population remained highly structured, with activity packets resembling those observed during wake. In contrast, LMN-HD population activity was less structured with respect to the cells’ preferred direction. Projections of population activity during non-REM sleep revealed a ring-like structure only for ADN, not for LMN (Figure 1H).

To quantify this phenomenon, we computed pairwise coupling (i.e., Pearson’s correlation of binned spike trains; see Methods) in both structures. As suggested by the low-dimensional projections, pairwise coupling among LMN-HD cells during sleep was less related to their coupling during wake (and thus, to their angular offset) than in the ADN (Figure 1I-J; Supp. Figure 2D-E).

Could the reduced structure in LMN during non-REM reflect a diminished ability to influence downstream ADN neurons? To address this, we measured LMN-ADN cross-correlograms at short time scales. These revealed strong, positively biased interactions from LMN to ADN consistent with synaptic transmission during both wakefulness and sleep (Figure 1K, Supp Figure 2F-G), suggesting that LMN maintains its capacity to drive ADN spiking.

Together, these observations argue against the presence of a stable attractor in the LMN during non-REM sleep and suggest instead that the ADN-HD population maintains coherent dynamics independently of its upstream input.

### During non-REM sleep, cortical feedback positively increases the coherence of LMN population activity

The decrease—but not complete abolition—of LMN-HD cell population coherence during non-REM could reflect intrinsic attractor-like properties of the LMN circuit and its inputs from the brainstem ^1,6,20,21^. Alternatively, this residual coordination may arise from cortical feedback, notably from the postsubiculum (PSB) ^18,19^. To test this possibility, we implanted *n* = 4 mice with probes targeting both the LMN and PSB (Figure 2A-D, Supp. Figure 3A-B). Cross-correlograms between spike trains in the two regions showed peaks at short time scales consistent with cortico-mammillary synaptic transmission (Figure 2E-F) for both wakefulness and sleep (Figure 2G).

**Figure 2.**
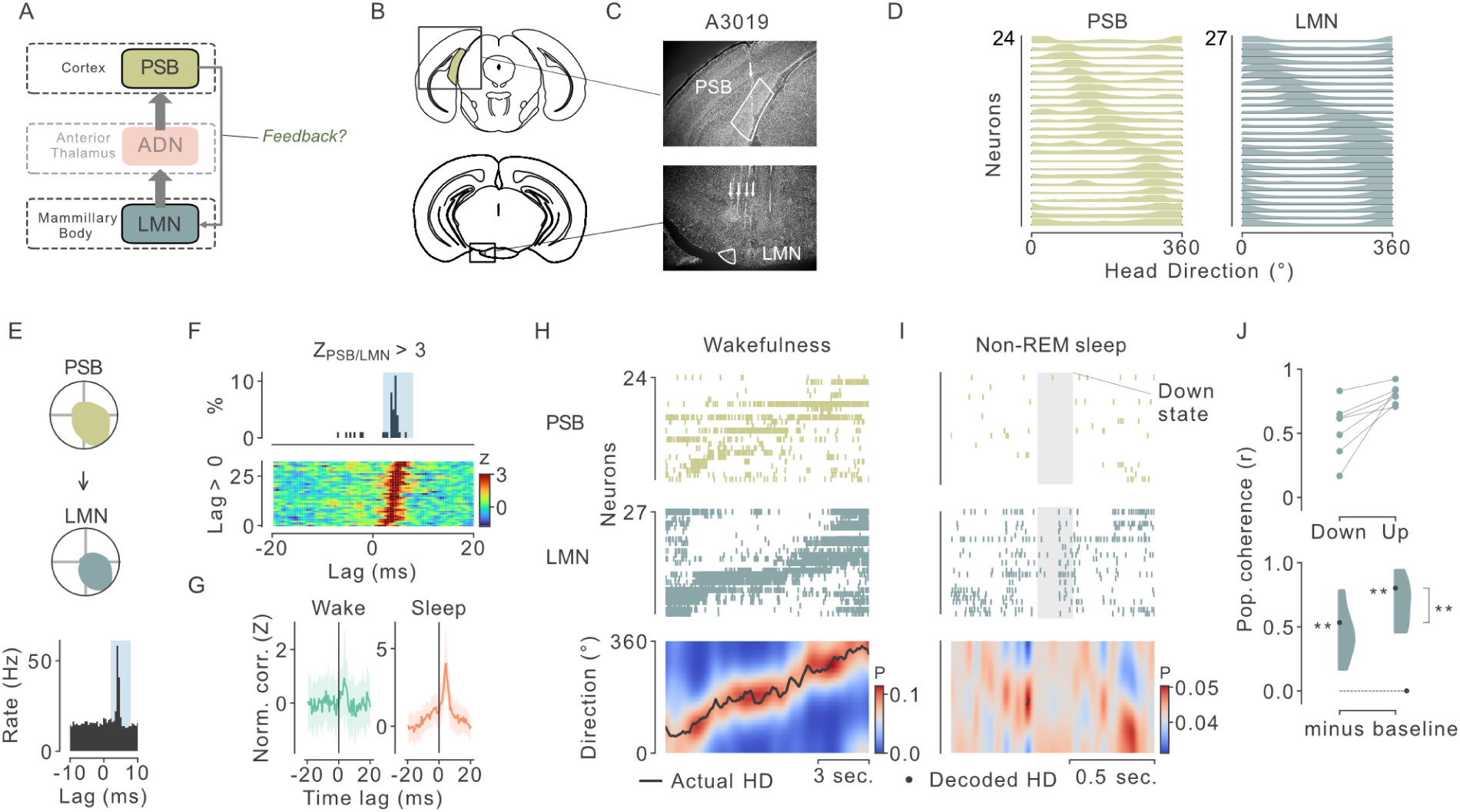
Reduction of LMN-HD cell activity coherence during cortical down states. **(A)** Schematic of the HD circuit emphasizing the cortical feedback from PSB to LMN. **(B)** Coronal diagrams showing LMN (bottom) and PSB (top) location. **(C**) Histological verification of probe locations (arrows) in the PSB (top) and LMN (bottom). **(D)** Tuning curves of HD neurons in the PSB (left) and LMN (right) sorted by preferred direction. **(E) T**op: tuning curve of two HD cells recorded simultaneously in the PSB and LMN. Bottom: cross-correlograms of the two cells spike trains showing a significant peak at short (4 ms) timelag. **(F)** Color-coded cross-correlograms (bottom) for all significantly coupled PSB→LMN pairs (Z > 3) and distribution of peak timelags (top). **(G)** Average normalized cross-correlograms of PSB→LMN cell pairs during wakefulness (green) and non-REM sleep (orange) show strong short-latency peaks consistent with monosynaptic influence from PSB to LMN in both states. **(H-I)** Simultaneous recordings of HD cells in PSB (yellow) and LMN (blue) during wakefulness (left) and non-REM sleep (right). Top: spike rasters. Bottom: actual (wake) or decoded (sleep) head direction. Gray area indicates a down state detected from PSB activity. **(J)** Top: LMN population coherence during cortical DOWN vs. UP states. Bottom: LMN coordination was larger during UP states compared to DOWN states, as measured by baseline-subtracted pairwise correlation (Wilcoxon-signed rank test; W=28; p<0.01; n=7 sessions). In both conditions, LMN coordination was larger than shuffle (Wilcoxon-signed rank test; W=28; p<0.01)

During non-REM sleep, thalamocortical networks display activity levels comparable to wakefulness during “UP” states, interrupted by brief periods of global silencing known as “DOWN” states ^22^ (Figure 2H-I). Crucially, LMN population coherence was decreased during cortical DOWN states (Figure 2I,J), indicating that PSB feedback may be necessary to organize LMN-HD activity.

### Optogenetically silencing cortical feedback disrupts the coherence of LMN population activity

To directly test the role of cortical feedback, we silenced PSB activity by transfecting local inhibitory neurons (using VGAT-Cre mice) with the red-light–activated depolarizing opsin Chrimson (Figure 3A, See Methods; ^23^). Red-light illumination of PSB led to robust, global suppression of PSB activity (Figure 3B, Supp. Figure 4A-D). Next, we injected Chrimson in the PSB of VGAT-Cre mice (*n* = 7 mice) and implanted probes in the ipsilateral LMN (Figure 3C, Supp. Figure 5), consistent with the lateralized nature of the PSB→LMN projection ^24–26^.

**Figure 3.**
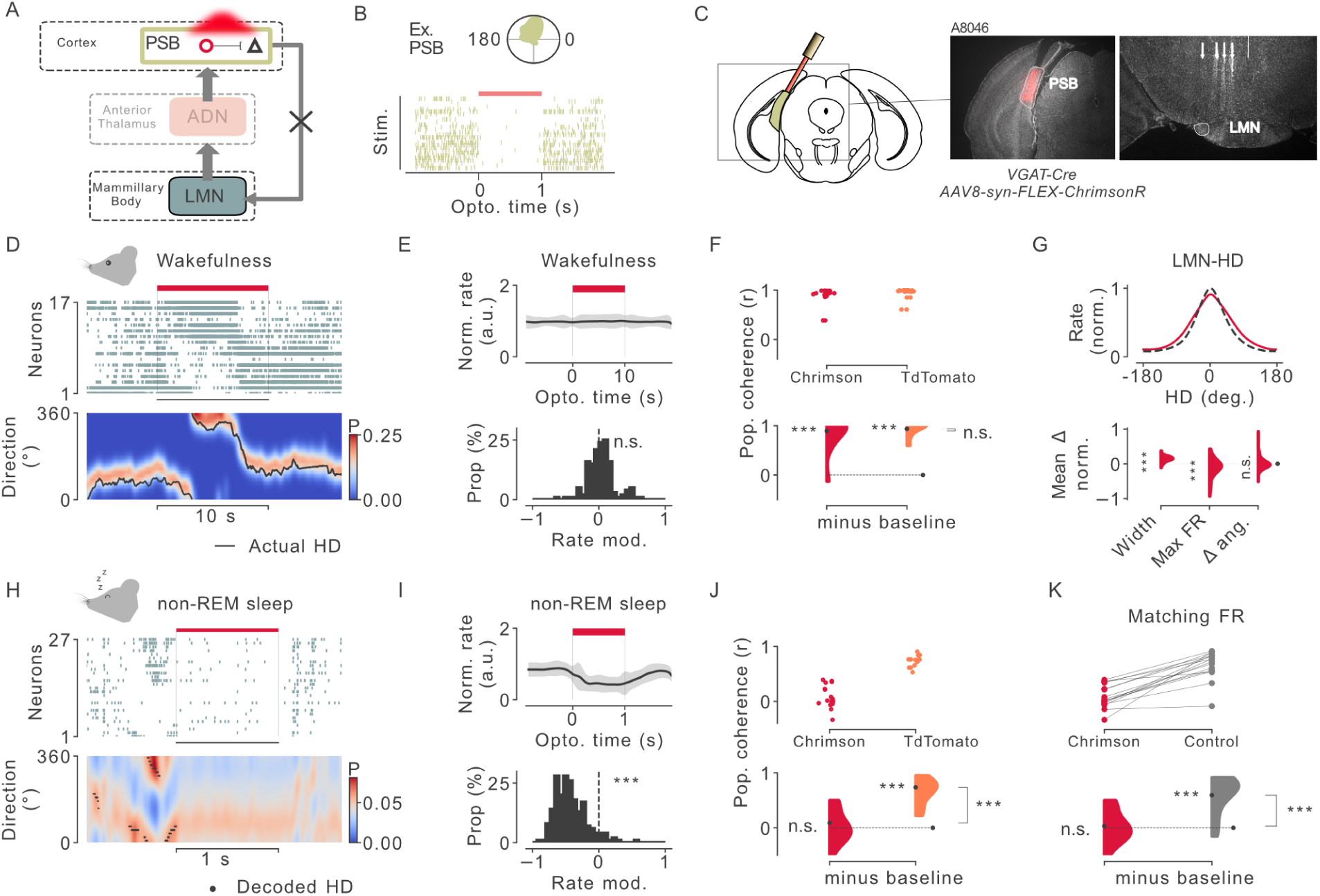
Cortical feedback is necessary for LMN-HD cell activity coherence during non-REM sleep. **(A)** Experimental design. VGAT-Cre mice received Cre-dependent Chrimson injections in PSB and were implanted with silicon probes in the LMN. **(B)** Example of a PSB-HD cell being inhibited during optogenetic stimulation of PSB VGAT-positive neurons. **(C)** Left: Coronal diagram of PSB showing optogenetic fiber placement. Right: Histological verification of PSB transfection, fiber location and shanks position for LMN recording. **(D)** Example wake session with PSB silencing (red bar). Top: LMN spike raster. Bottom: decoded HD signal in the LMN remains coherent. **(E)** Top: Normalized LMN firing rate during PSB silencing in wakefulness. Bottom: Distribution of modulation indices shows no significant change (Wilcoxon-signed; W=4465; p=0.10; n=145 neurons). **(F)** LMN population coherence during wake in Chrimson (red) and control (TdTomato, orange) mice. No significant difference across conditions (Mann-Whitney; U=188; p=0.22; n_Chrimson_=24; n_TdTomato_=20), and coherence remained higher than chance in both (U_Chrimson_=572; p < 0.001; U_TdTomato_=400; p<0.001). **(G)** Top, Average centered LMN-HD cell tuning curves during control periods (dashed line) and PSB silencing (red). Bottom, Change in tuning curve properties (width, peak firing rate, and preferred direction) during PSB silencing in wakefulness (Wilcoxon-signed; W_Widths_=107; p<0.001; W_Max FR_=1376; p<0.001; W_ΔAng_=4187; p=0.19). **(H)** Same as (G) during a non-REM sleep session. **(I)** Top: Change in mean LMN firing rate during PSB silencing in non-REM sleep. Bottom: Distribution of modulation indices (Wilcoxon-signed; W=349; p<0.001; n=204 neurons). **(J)** Change in LMN-HD cell population coherence during non-REM sleep when PSB was silenced compared to TdTomato control mice (Mann-Whitney; U=11; p<0.001; n_Chrimson_=16; n_TdTomato_=15). Coherence was not different from chance with Chrimson (Mann-Whitney; U=136; p=0.38) and was unaffected in TdTomato mice (Mann-Whitney; U=225; p<0.001) **(K)** Difference in coherence after matching LMN-HD cell firing rates between Chrimson and control mice, ruling out that change in HD cell coherence resulted from decreased firing rates (Mann-Whitney; U_across_=28, p < 0.001; U_Chrimson_=136; p=0.38; U_TdTomato_=236; p<0.001; n=16 sessions).

During wakefulness, silencing PSB did not significantly affect LMN firing rates or population coherence (Figure 3D-G). However, tuning curves of LMN-HD cells became slightly broader (Figure 3G), which would indicate unstable or reduced precision tuning, consistent with a role for PSB feedback in anchoring the internal (and still coherent) HD signal to external environmental cues and landmarks ^16,24^.

By contrast, silencing PSB during non-REM induced profound changes in LMN population activity. LMN firing rates dropped to approximately half of their baseline levels (Figure 3H-I), and—critically—population coherence became indistinguishable from random spiking (Figure 3J-K).

Together, these results reveal a crucial role for cortical feedback in organizing coherent population dynamics in the LMN-HD system during non-REM sleep.

### A coherent HD signal is generated in the ADN during non-REM

The previous findings suggest that the ADN–PSB network can sustain structured population activity even when its upstream input from the LMN becomes disorganized during non-REM. However, since the PSB projects back to the thalamus both directly and indirectly ^27,28^, it remains possible that the ADN still depends on cortical feedback to maintain a coherent HD signal. Alternatively, coherent dynamics may arise intrinsically within the thalamus itself.

To test this, we repeated the PSB silencing experiment while recording from the ADN-HD cell population (*n* =7 mice; Figure 4A–B; Supp Figure 6). During wakefulness, ADN-HD cell firing rates were reduced by ∼17% only for bilateral silencing of PSB (Figure 4C-D, G). We observed no significant differences in ADN population coherence for both ipsilateral and bilateral PSB silencing during wakefulness (Figure 4H).

**Figure 4.**
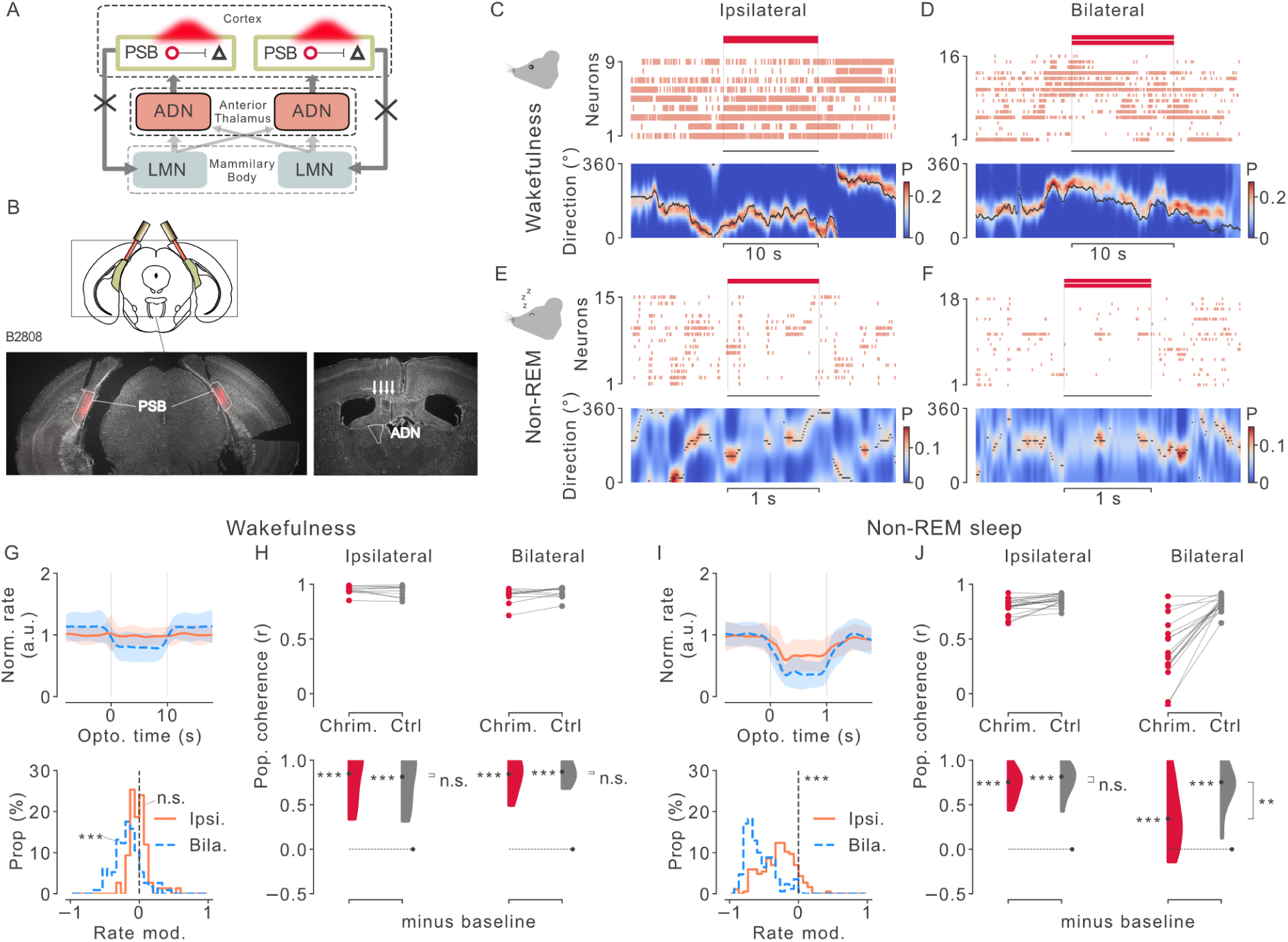
ADN-HD cells maintain coherent population activity during cortical silencing across brain states. **(A)** Schematic of cross-hemispheric HD system circuit. ADN receives input from both the ipsilateral and contralateral LMN. **(B)** Experimental design. Top: VGAT-Cre mice are transfected with Cre-dependent Chrimson AAV in PSB bilaterally, and were implanted with silicon probes in the ADN. Bottom: Histological verification of PSB transfection and fiber location for bilateral injections and ADN targeting. **(C)** Example of ADN-HD population activity during wakefulness with unilateral PSB silencing (red bar). Top: spike raster. Bottom: decoded head direction. **(D)** Same as (C) with bilateral silencing of the PSB. (E-F) Same as (C-D) during non-REM sleep. **(G)** Top: Normalized ADN firing rate during ipsilateral (red) and bilateral (blue) PSB silencing as animals were awake. Bottom: Histogram of modulation indices for ipsilateral (Wilcoxon-signed; W=1082; p = 0.07; n=75 neurons) and bilateral (W=620; p<0.001; n=114 neurons; ∼17% decrease). **(H)** Top: Wake ADN-HD population coherence during ipsilateral (left) and bilateral (right) PSB silencing compared to TdTomato controls (using matched firing rates). Bottom: baseline-subtracted coherence for ipsilateral (Mann-Whitney; U_Across_=55; p=0.73; U=100; p < 0.001 for Chrimson & control compared to baseline) and bilateral (U_Across_=49; p=0.96; U=100; p < 0.001 for Chrimson & control). **(I-J)** Same as (G-H) during non-REM sleep. (I) change in firing rates for ipsilateral (Wilcoxon-signed; W=492; p < 0.001; n=186 neurons; ∼30% decrease) and bilateral conditions (W=6; p<0.001; n=170 neurons; ∼56% decrease). (J) ADN-HD population coherence for ipsilateral (Mann-Whitney; U_Across_=143; p=0.28; U=361; p<0.001 for both conditions compared to baseline) and bilateral conditions (U_Across_=50; p<0.01; U_Chrimson_=240; p < 0.001; U_Control_=286; p<0.001).

During non-REM sleep, ADN firing rates were reduced by ∼30% for unilateral silencing of the PSB and ∼56% decrease during bilateral silencing (Figure 4E-F, I), likely reflecting the overall reduction of excitatory inputs from bilateral LMN. Strikingly, however, the ADN-HD population maintained internal coherence at levels indistinguishable from controls (Figure 4J) for unilateral silencing of PSB. This resilience may reflect the fact that, unlike the ipsilateral PSB→LMN projection, the ADN receives input from both left and right LMN ^6^. During bilateral PSB silencing, the coherence of the ADN-HD population decreased, yet remained higher than control conditions (Figure 4J). Together, these results indicate that coherent HD dynamics can emerge locally within the anterior thalamus during non-REM sleep, independently of structured cortical or subcortical input.

### Evidence for non-linear input–output transformations in ADN-HD neurons

Mechanistic models of attractor networks typically rely on strong local excitation combined with broad inhibitory connectivity, forming a structure akin to a passband filter in feature space that supports the emergence of a coherent activity packet ^21,30^. However, thalamocortical relay cells lack local excitatory collaterals ^31^. Instead, they interact primarily via shared connections to inhibitory neurons in the thalamic reticular nucleus (TRN) ^7,26,27^. It is unclear how thalamic circuitry could support the emergence of structured activity.

Building on previous work ^29^, we observed that ADN-HD cells do not fire in a smooth, graded manner as the head moves through their tuning field. Rather, they show abrupt transitions from a low-firing “ground state” to a high-firing “activated state” (Figure 5A–B) ^29^. This non-linearity was reflected in the interspike interval distributions, which were more bimodally distributed in ADN-HD cells compared to LMN-HD cells (Figure 5C-D).

**Figure 5.**
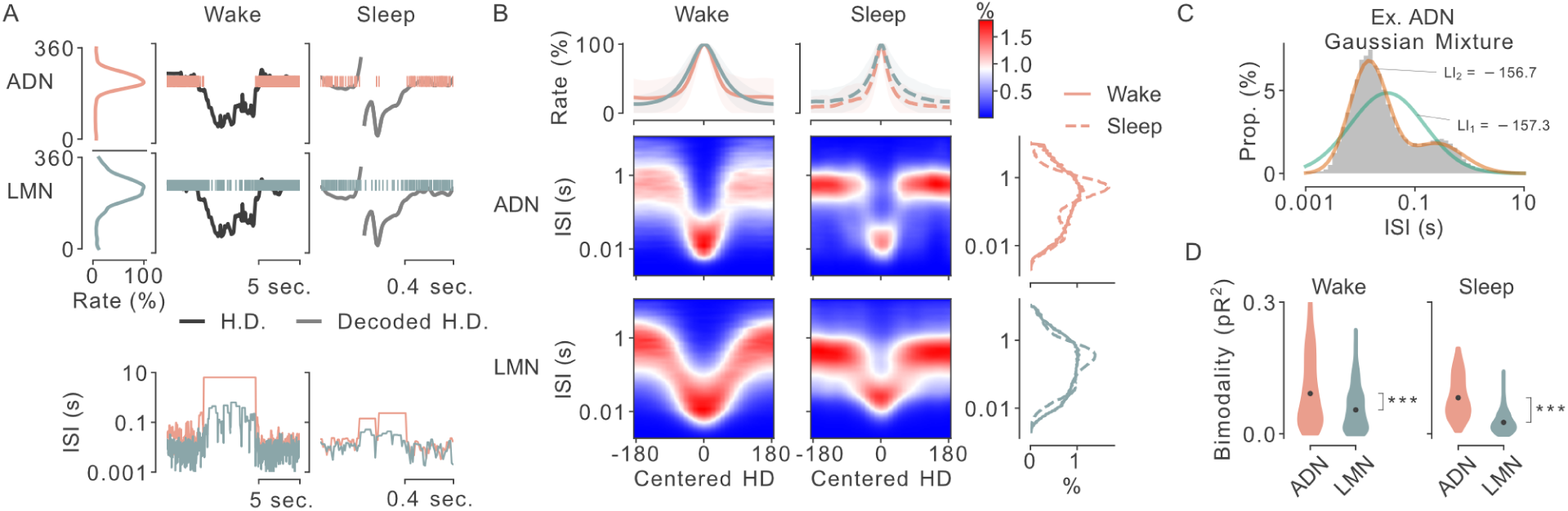
Strong non-linear responses of ADN-HD cell population. **(A)** Example activity of two ADN and LMN-HD cells during wake and non-REM sleep. Top Left: tuning curves. Top Middle: head direction (black) and firing activity (colored) showing transitions between ground and active states. Top Right: decoded HD signal during non-REM and spiking activity. Bottom: corresponding interspike intervals (ISI) during wakefulness (left) and non-REM sleep (right). **(B)** Average firing rate (top) and ISI distributions (middle) as a function of centered HD angle, aligned to each cell’s preferred direction, for ADN (top row) and LMN (bottom row) during wake (left) and sleep (right). Right: marginal ISI distributions. **(C)** Example ADN-HD cell fit with a two-component Gaussian mixture model, indicating bimodality. Bottom: quantification of bimodality index (pseudo-R²) across cells in ADN and LMN during wake (Mann-Whitney; U=57232; p<0.001) and sleep (U=78931; p < 0.001).

### Circuit non-linearity and inhibition are sufficient to generate ADN HD structure

Could such strong non-linearity be sufficient to explain the generative dynamics observed in the ADN? To address this, we simulated a simplified circuit comprising the three principal stages of the HD system studied here—LMN, ADN, and PSB—connected in a closed loop (Figure 6A, Supp Figure 7A-B). The model incorporated three key features: (i) the LMN→ADN projection is divergent, with each LMN neuron contacting multiple ADN targets (as the number of neurons in the ADN is greater than in the LMN); (ii) the ADN is reciprocally connected with a local inhibitory population; and (iii) ADN neurons exhibit strong input–output non-linearity.

**Figure 6.**
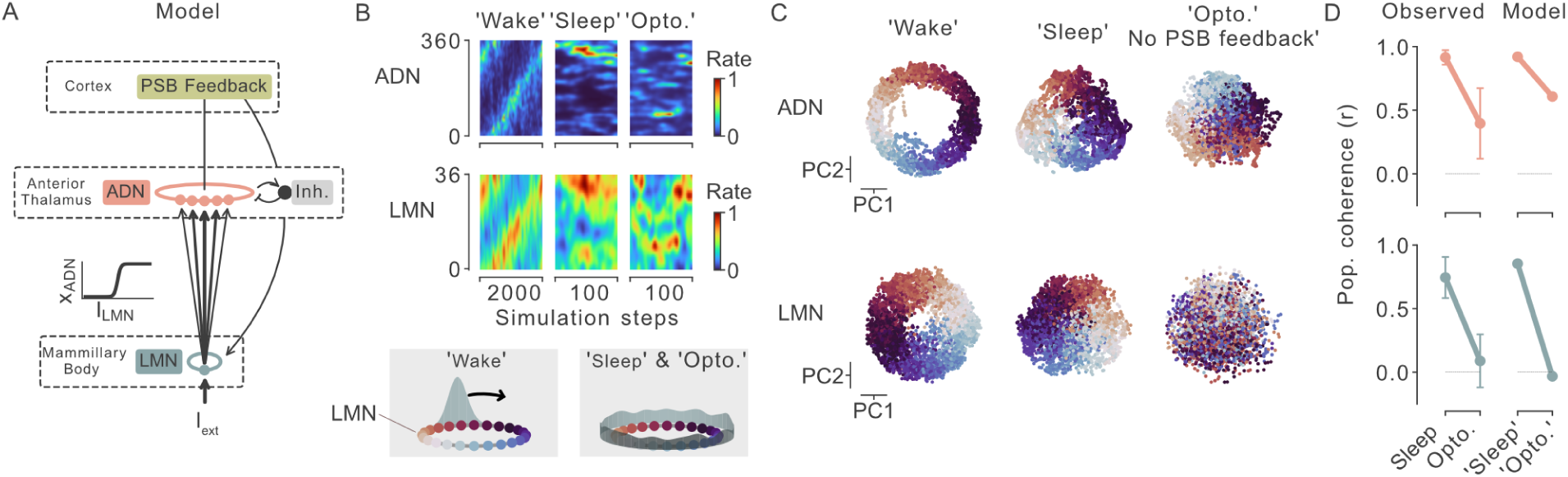
Strong non-linear responses and global inhibition are sufficient to account for the generative properties of the ADN-HD cell population. **(A)** Schematic of the computational model incorporating LMN→ADN divergence, ADN inhibition, and PSB feedback. ADN neurons exhibit threshold-like nonlinear responses. **(B)** Population activity of simulated HD cells in ADN (top) and LMN (bottom) under three conditions: ’Wake’-like input, ’Sleep’-like input (i.e. random inputs), and PSB silencing during ‘sleep’ (’Opto.’). **(C)** Projections of population activity for the three conditions for LMN (bottom) and ADN (top) showing the preserved ring structure for ADN. **(D)** Population coherence values from actual data (left) and the model (right), comparing sleep and optogenetic PSB silencing, for ADN (top) and LMN (bottom).

In the closed-loop configuration, with LMN neurons driven by noise and feedback from PSB to LMN intact, the circuit spontaneously generated a coherent activity packet (Figure 6B-C). As expected, when the loop was opened (i.e., PSB→LMN feedback removed), LMN activity remained unstructured. Remarkably, however, the ADN-HD population still exhibited significantly greater structure than predicted by chance (Figure 6C-D, Supp Figure 7 C). These simulations support the idea that structured HD dynamics can emerge in the anterior thalamus from random input, provided sufficient non-linearity and inhibitory shaping within the circuit.

## Discussion

In this study, we performed multi-site electrophysiological recordings and optogenetic manipulations *in vivo* to reveal how the thalamus supports coherent population activity in the head-direction system. Specifically, we show that during non-REM sleep, the pairwise correlation of LMN-HD cells is lower than during wakefulness, unlike ADN-HD cells, which preserve high coherence across brain states. Transient inactivation of cortical feedback from the PSB with optogenetics led to a total decorrelation of LMN-HD cells, while maintaining coherent dynamics of ADN-HD cells. This indicated that thalamic circuits are capable of generating a coherent HD signal irrespective of the coherence of their inputs.

The data and simulations support the view that coherence in the ADN can emerge *de novo* from noise, provided the circuit architecture includes divergent input, local inhibitory interactions, and sharp non-linear response thresholds. This mechanism is particularly compelling in the thalamus, where recurrent excitation is anatomically absent, and suggests a more active, generative role for thalamic nuclei than traditionally assumed.

The generation of the HD signal during wakefulness is thought to rely on reciprocal interactions between the dorsolateral tegmentum and the LMN ^5,6,32,32,33^. Our findings suggest that this circuit becomes inoperative during non-REM sleep, with the weak coordination observed among LMN neurons instead being driven by cortical feedback. Notably, the structured activity of the LMN re-emerges during REM sleep, pointing to a state-dependent gating of internal dynamics. Neuromodulatory changes—particularly in cholinergic tone—may play a central role in this transition ^34,35^.

In our model, coordinated activity among ADN-HD neurons requires local & non-specific inhibitory regulation. This inhibition can originate from the TRN, which is reciprocally connected with most thalamic nuclei ^30^, including the ADN ^7,27,36^. Whether TRN neurons provide only broad, non-specific inhibition or contribute more selectively to the organization of HD activity—potentially through structured interactions among TRN neurons themselves—remains an unresolved question for future investigation.

For much of modern neuroscience, the thalamus has been regarded primarily as a relay hub: a subcortical structure that gates and modulates the flow of sensory information to the cortex ^37,38^. Yet, recent studies have begun to reveal that thalamic circuits may participate in more complex operations—including state-dependent modulation, attention, and memory-related processes ^39–44^. Our findings suggest that the thalamus is endowed with generative capabilities, supporting structured population dynamics in the absence of external input. The spontaneous and structured activity of HD cells during sleep is thought to coordinate activity in downstream structures such as the medial entorhinal cortex ^4,45,46^ and the hippocampus ^47^, and may thereby contribute to memory consolidation and replay processes ^11,48–50^.

Altogether, these findings support the view that during non-REM sleep, thalamocortical networks disengage from external input and shift toward intrinsically generated population dynamics ^51^.

## MATERIALS AND METHODS

### Animals

All experiments were approved by the Animal Care Committee of the Montreal Neurological Institute. Adult (>8 weeks old) male C57BL/6J and VGAT-IRES-Cre (homozygous and heterozygous) mice were used. C57BL6/6J and homozygous VGAT-IRES-Cre animals were obtained from Jackson Laboratories (000664 and 028862 respectively), while heterozygous VGAT-IRES-Cre animals were bred by crossing homozygous VGAT-IRES-Cre males with C57BL/6J females. Animals were given food and water *ad libitum* and kept under a 12hr light/dark cycle.

### Silicon probe implantation

Animals were anesthetized with isoflurane (5% at induction, 1-2% at maintenance, airflow 2 L min^-1^) and subsequently placed in a stereotaxic frame (Kopf Instruments). Animals were injected subcutaneously with carprofen (20 mg kg^-1^ body weight) and kept on a heating pad for the entire procedure. Eyes were covered with lubricant eye gel (Systane) before removing fur with Veet hair removal cream. The skin was disinfected with 2% Chlorhexidine before applying 5% lidocaine cream. Surgical instruments were sterilized with a Geminator 500 before opening the skin with a surgical blade. A silver wire was inserted into the cerebellum to serve as a reference and ground. All probes were mounted on a moveable microdrive. Three types of probes were used according to the target region: long four-shank probes (Neuronexus Buzsaki32L) were used to target LMN, short four-shank probes (Neuronexus Buzsaki32S) were used to target ADN, and single-shank probes (Cambridge NeuroTech H5) were used to target PSB. A hole was drilled into the skull above the target site large enough to accommodate the size of the probe. Four-shank probes were implanted perpendicular to the midline. Probes were slowly lowered above the target site and secured to the skull using a light-cure adhesive (Kerr OptiBond) and dental acrylic cement (Unifast Trad). A copper mesh was attached with a flowable composite (FusionFlo) around the probe to reduce electrical noise in the recordings. Following the implant, Animals were injected subcutaneously with saline (1 mL) and carprofen (20 mg kg^-1^) and placed on a heating pad until normal behaviour resumed. Carprofen injections were delivered every 24 hours for 72 hours after implantation. All animals were housed individually following the implant surgery and allowed to recover for at least one week before electrophysiological recordings. Silicon probe implantation coordinates were as follows: LMN implant AP (from Bregma): -2.23, ML (from superior sagittal sinus): -0.75, DV (from dura): -4.90; ADN implant AP (from Bregma): -0.4, ML (from superior sagittal sinus): -0.72, DV (from dura): -2.32; PSB implant AP (from Lambda): +0.30, ML (from superior sagittal sinus): -2.35, DV (from dura): -1.00.

### Optogenetic experiments

Animals were injected with either the viral vector AAV8-Syn-Flex-ChrimsonR-tdT (UNC; batch number AV5844D) or AAV8-CAG-Flex-tdTomato (UNC; batch number AV4912b), with concentrations of 3.9×10^12^ and 5.5×10^12^ respectively. Viruses were subdivided into aliquots and stored at -80 °C until use.

Animals were anesthetized using isoflurane and placed on the stereotaxic frame as previously described. After exposing the skull, a small hole was drilled over the target site. Viruses were injected at a speed of 50 nl min^-1^ for a total volume of 100 nL into the target site using a Nanofil syringe (WPI) and a Pump 11 Elite Nanomite Programmable Syringe Pump (Harvard Apparatus) mounted on the stereotaxic frame. Following injection, the needle was left in place for 10 minutes to allow for proper diffusion and absorption of the virus before being slowly removed from the brain. Unilaterally injected animals were injected in the left hemisphere at a 30° angle towards the midline with the following coordinates: AP (from lambda): +0.28 mm, ML (from superior sagittal sinus): -1.21, DV (from dura): -2.20. Bilaterally injected animals were injected at a 28° angle towards the midline with the following coordinates: AP (from lambda) +0.28, ML (from superior sagittal sinus) ± 1.33, DV (from dura) -1.90.

Animals were sutured (Ethicon Perma-hand Silk) and injected subcutaneously with sterile saline (1 mL) before being placed back in their home cage over a heating pad until normal behaviour resumed (max 30 mins). Animals were subcutaneously injected with carprofen every 24 hours for 72 hours following the procedure and were allowed a recovery period of at least 14 days before the implantation surgery.

Optic fibers (Doric Lenses; MFC_200/240-0.22_27mm_RM2_FLT) were implanted unilaterally (left hemisphere) at a 30° angle towards the midline with the following coordinates: AP (from lambda) +0.30, ML (from superior sagittal sinus) -1.10, DV (from dura) -1.50. All unilaterally implanted ADN animals were also implanted with tungsten wires targeting the pyramidal layer of CA1. Optic fibers were implanted bilaterally at a 28° angle towards the midline with the following coordinates: AP (from lambda) +0.28 or +0.52, ML (from superior sagittal sinus) ± 1.23, DV (from dura) -1.45. Fibers were secured to the skull using a combination of a flowable composite (FusionFlo) and dental acrylic cement (Unifast Trad). Light power output of optic fibers was measured before each implantation and an output curve was calculated for each fiber. Light output was set to 15mW for each recording session and laser light was delivered from a 638 nm fiber-coupled laser diode module (Doric Lenses), controlled with a laser diode module driver (Doric Lenses), to the optic fiber implants via patch cords. During sleep recordings, light pulses (1s) were delivered at 0.2 Hz for 5 mins, followed by 5 mins of no stimulation. This cycle was repeated four times for a total of 240 light pulses in a 45-min recording session. During wake recordings, light pulses (10s) were delivered at 0.5 Hz for 5 mins, followed by 5 mins of no stimulation. This cycle was repeated four times for a total of 60 light pulses over a 45-min recording session.

### Recording procedure

Animals were recorded during random foraging behaviour either in a square (80 cm width, 50 cm height) or a circular (85 cm diameter, 50 cm height) open field. Both recording chambers consisted of a metal frame supporting a gray plastic platform with gray walls. A white rectangular cue card served as a salient landmark in the circular environment, while the square environment contained different rectangular cue cards on three walls. Sleep sessions were performed in the home cage.

Neurophysiological signals were acquired continuously at 20 kHz on a 256-channel RHD USB interface board (Intan Technologies) and captured with Intan RHX software (Intan Technologies). Recording cables were tethered to a motorized electrical rotary joint (AERJ; Doric Lenses). Probes were lowered into the target structure in small (∼35 μm) increments. Data collection began at least 2 hours after the last adjustment.

The animals’ position and orientation were tracked in 3D using several infrared cameras (Flex 13; Optotrack) located above the enclosure and coupled to the Motive motion capture system (Optitrack). Position and head orientation were sampled at 120 Hz and synchronized with the electrophysiological recording via voltage pulses registered by the RHD USB interface board (Intan Technologies). Tracking markers were attached to the headcap. Video recording was captured by an overhead camera (Flex 13; Optitrack) placed close to the rotary joint.

### Tissue processing

Following the termination of experiments, animals were deeply anesthetized using a sodium pentobarbital solution injected intraperitoneally and perfused transcardially with a 0.9% phosphate-buffered saline solution followed by 4% paraformaldehyde. Brains were extracted and stored in 4% paraformaldehyde for 12-24 hours before being transferred to a 30% sucrose solution until the brains sank (48-72 hours). Brains were subsequently frozen to -80 °C and sectioned with a freezing microtome coronally in 40 μm slices. Sections were washed, counterstained with DAPI and mounted on glass slides with ProlongGold fluorescence antifade medium. Sections containing probe tracts, with the exception of optogenetic animals injected and implanted in the PSB, were additionally incubated with a Cy3 anti-mouse secondary antibody (1:200 dilution; Cedarlane, 715-165-150) to help visualize the electrode tract. A widefield fluorescence microscope (Leica) was used to obtain images of sections and verify the tracks of silicone probe shanks, optic fiber position and viral expression.

### Spike sorting and unit classification

Spike sorting was performed semi-automatically using Kilosort 4.0 ^52^ followed by manual curation using the software Klusters ^53^. At this stage, clusters lacking a distinct waveform and a clear refractory period in their autocorrelogram (0–1 ms bin) were labeled as noise. Cluster pairs showing similar waveforms and a clear refractory dip in their cross-correlograms (0–1 ms bin) were merged.

### Sleep State classification

To perform sleep scoring, tungsten wires were inserted in the pyramidal layer of CA1 in 7 animals and sleep stages were determined according to time-resolved spectrograms of the CA1 and animals’ movements. For the animals without CA1 wires, sleep scoring was performed by detecting theta frequency in the local field potential and animal movement. In all cases, stages were visually inspected and manually adjusted.

### HD tuning curves and selection

HD tuning curves were computed by dividing the spike count histogram by the occupancy time histogram, using 3° bins, and smoothing the result with a Gaussian kernel with a standard deviation of 12°. For each cell, HD information was computed as:

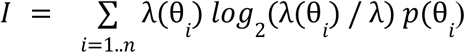

where λ(θ_*i*_) denotes the firing rate of the cell in the i-th angular bin, λ is the cell’s overall average firing rate during exploration, and *p*(θ_*i*_) represents the occupancy probability—that is, the normalized time spent facing direction θ_*i*_. The resulting information rate (in bits per second) was then divided by the cell’s mean firing rate to yield the information content (in bits per spike), providing a measure independent of firing rate.

HD cells were identified through a two-step selection process. First, cells with a mean firing rate greater than 1 Hz and HD information exceeding 0.1 were retained. In the second step, the exploration epoch was divided into two equal halves, and tuning curves were computed separately for each segment. A Rayleigh test for directional tuning was applied, and cells with a Rayleigh score above 50 and a p-value below 0.0001 were selected. Only cells that met these criteria in both sub-epochs were classified as HD cells.

### Decoding head-direction from population activity

Head-direction was decoded from spiking activity using a Bayesian approach implemented in the *pynapple* package ^54,55^. For the decoding shown in Figures 1–3 and Supplementary Figure 7, spike counts were computed in 100 ms bins during wakefulness and 5 ms bins during non-REM sleep. Tuning curves were calculated over 24 angular bins spanning 0° to 360°. For sleep decoding, only angles decoded from a non-uniform probability distribution are shown. Non-uniformity was assessed at each time step by thresholding the probability distribution, retaining only those with a normalized entropy less than 0.12.

### Dimensionality reduction

Low-dimensional projections of population activity were computed using kernel principal component analysis (kPCA). Spike trains were discretized into non-overlapping time bins of 300 ms during wakefulness and 20 ms during non-REM sleep, reflecting the distinct temporal dynamics of these states. To stabilize variance across neurons and reduce the influence of high firing rates, spike counts in each bin were square-root transformed. The resulting time series were then temporally smoothed by convolution with a Gaussian kernel whose standard deviation was set to three times the bin width. The preprocessed population activity vectors were subsequently embedded into a low-dimensional space using kernel PCA with a cosine similarity kernel, which emphasizes relative firing patterns across neurons while being insensitive to global fluctuations in population rate. Projections were computed using the first two principal components.

### Population coherence

Population coherence was computed as the correlation between the correlation coefficients of pairs of simultaneously recorded neurons. Spike trains were binned using 100 ms windows for wakefulness and REM sleep, and 20 ms windows for non-REM sleep. The resulting spike count vectors were convolved with a Gaussian kernel with a standard deviation equal to three times the bin width. Baseline population coherence was obtained by shuffling neuron identities.

To match firing rates between conditions during optogenetic stimulation, we first computed the change in firing rate between the stimulation epoch and the immediately preceding control epoch. If the change was negative, spikes from the control epoch were randomly subsampled to match the firing rate observed during stimulation. This subsampling procedure was repeated 1,000 times, and the Pearson correlation for each neuron pair was averaged across all samples.

### Cross-correlograms

Cross-correlograms between pairs of neurons were computed using 100 ms bins for wakefulness and 10 ms bins for non-REM sleep.

### Detection of monosynaptic connections

Spike train cross-correlograms were computed over a ±100 ms window using 0.5 ms bins and convolved with a Gaussian kernel (σ = 10 ms) to estimate the baseline firing rate. The raw cross-correlograms were then z-scored relative to this baseline. A putative monosynaptic connection was deemed significant if at least two consecutive bins within the 2–8 ms interval exceeded 3 standard deviations above the baseline.

### Detection of Up and Down states

Down states were detected by pooling spikes from all PSB units. The combined spike trains were binned in 10 ms intervals and smoothed with a Gaussian kernel whose standard deviation was twice the bin size. Down states were identified by thresholding at the 50th percentile of the smoothed firing rate during non-REM sleep. Down epochs separated by less than 100 ms were merged, and epochs shorter than 20 ms or longer than 2 s were excluded. Up states were defined as the complement of down states during non-REM sleep.

### Gaussian mixture of ISI distribution

Gaussian mixture models were fitted using the *scikit-learn* package. Only ISI values between 2 ms and 10 s were used to fit the Gaussian models. Bimodality was quantified using the pseudo-R² metric: 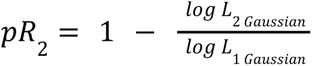

### Network model

We implemented a recurrent neural network model comprising three interconnected populations representing the lateral mammillary nucleus (LMN), anterior dorsal thalamic nucleus (ADN), and thalamic reticular nucleus (TRN). The model was implemented in Python using NumPy and Numba for computational efficiency.

The network consisted of *N*_*LMN*_ = 36 units, *N*_*ADN*_ = 360 units and a single unit representing the TRN. All units were modeled as rate-based neurons with first-order dynamics governed by the differential equation:

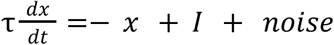

where x is the membrane potential variable, τ=0.1 is the integration time constant, I represents synaptic inputs and external drives, and noise is an additive Gaussian term modeling stochastic fluctuations.

LMN and TRN activity is governed by linear thresholding (rectified linear units) with *r*_*LMN*_ = *max*(0, *x*_*LMN*_) and *r*_*TRN*_ = *max*(0, *x*_*TRN*_), whereas ADN units used a sigmoidal transfer function:

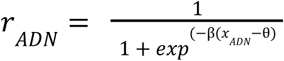

with slope parameter β=3 and threshold θ=1.

Independent Gaussian noise was injected into each population to capture intrinsic neuronal variability, with standard deviations parameterized as σ_*LMN*_ = 1, σ*_ADN_* = 0. 1 and σ_*TRN*_ = 0. 1

During wakefulness, LMN units received a rotating external input, implemented as a periodically shifted Gaussian bump, mimicking a HD signal. A constant drive term *D*_*LMN*_ = 1 was added to LMN units to maintain baseline activation. An external input scaling parameter *I*_*LMN*_ controlled the strength of the external input drive, which was set to 0.43 during wakefulness and set to zero to simulate sleep.

Connections from LMN to ADN were arranged in a circular topology using Gaussian-weighted synaptic weights with standard deviation σ=100 (Supp. Figure 7). This circular connectivity ensured that ADN units received inputs preferentially from LMN units coding nearby directions on the ring.

The TRN unit received excitatory input from the summed activity of ADN neurons, scaled by *w*_*ADN* → *TRN*_ = 1. 0. In turn, the TRN inhibited ADN neurons through feedback weights scaled by *w*_*TRN* → *ADN*_ = 0. 05. This inhibitory feedback loop controlled the excitability of the ADN population and regulated its dynamics.

A feedback pathway from ADN to LMN was implemented to simulate the feedback from PSB. This connection used a circular weight matrix with similar tuning (σ=100). PSB feedback matrix was scaled by a factor *w*_*PSB* → *LMN*_ = 0. 11.

During “wakefulness”, external input to LMN was enabled and feedback pathways were active. During “sleep”, external input to LMN was suppressed. During “optogenetic”, feedback weights from the ADN units back to the LMN units were artificially set to 0.

### Data analysis and statistics

All analyses were done in python with the following libraries: Scipy, Numpy, Matplotlib, Scikit-Learn and Pynapple ^55^.

## AUTHOR CONTRIBUTIONS

Conceptualization: GV, SSC, AP; Methodology: GV, SSC, AP; Software: GV, AP; Validation: GV, SSC, AP; Formal analysis: GV; Investigation: GV, SSC; Resources: AP; Data Curation: GV, SSC; Writing - Original Draft: GV, SSC, AP; Writing - Review & Editing: GV, SSC, AP; Visualization: GV, SSC, AP; Supervision: AP; Project administration: AP; Funding acquisition: AP

## ACKNOWLEDGMENTS

We would like to thank Lynda Mainville for technical support. We are thankful to the members of Peyrache laboratory for comments on the earlier version of the manuscript. This work was supported by a Canadian Research Chair in Systems Neuroscience (AP), CIHR Project Grants 155957, 180330, and 190289 (AP), NSERC Discovery Grants RGPIN-2018-04600 and RGPIN-2025-06801 (AP), and Vanier CGS (SSC).

## SUPPLEMENTARY INFORMATION

### Supplementary figures 1–8

**Supplementary figure 1.**
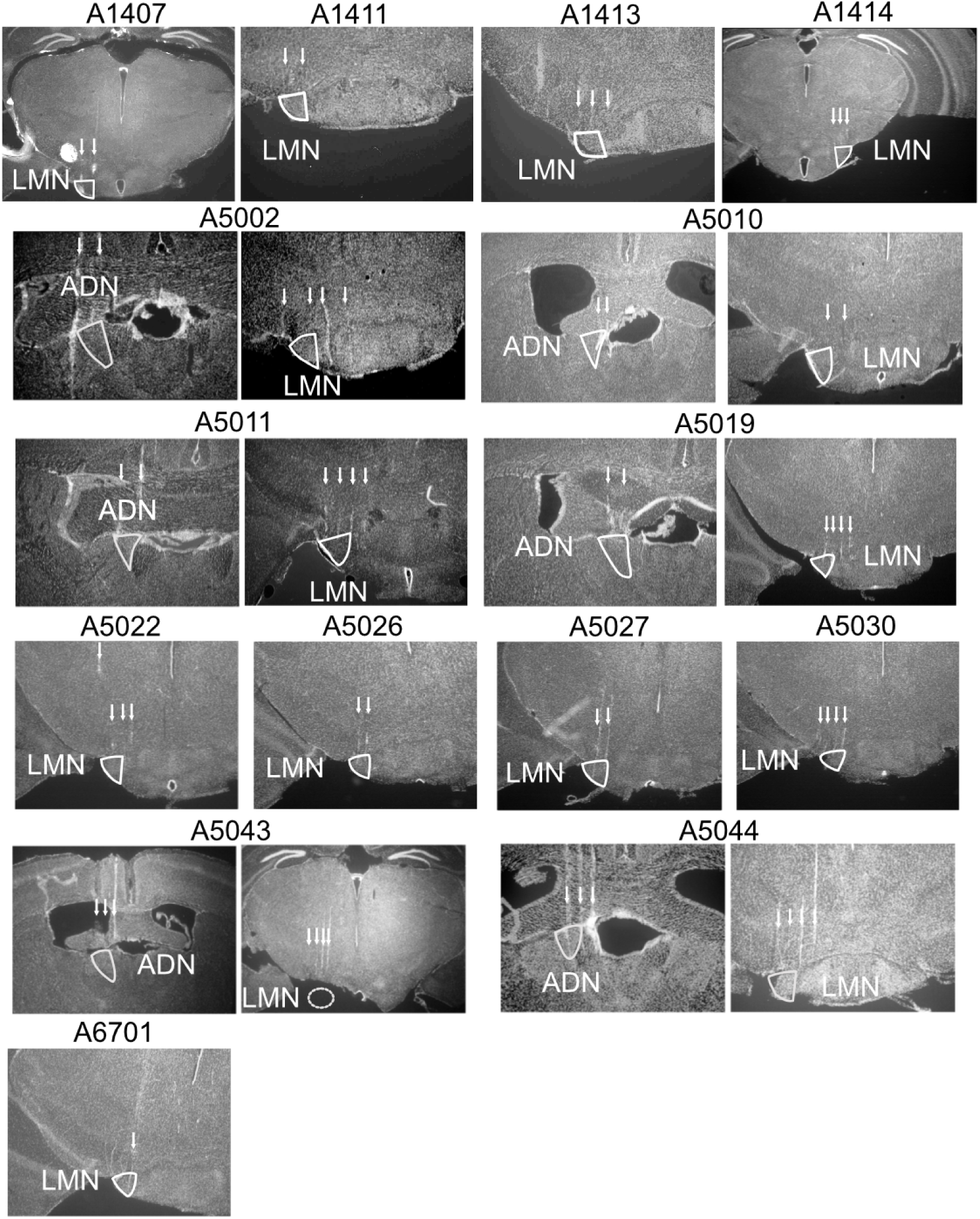
Histological validation of electrode placement in the ADN and LMN. Coronal histology showing silicon probe tracks (arrows) targeting the LMN and/or ADN for all animals presented in Figure 1.

**Supplementary Figure 2.**
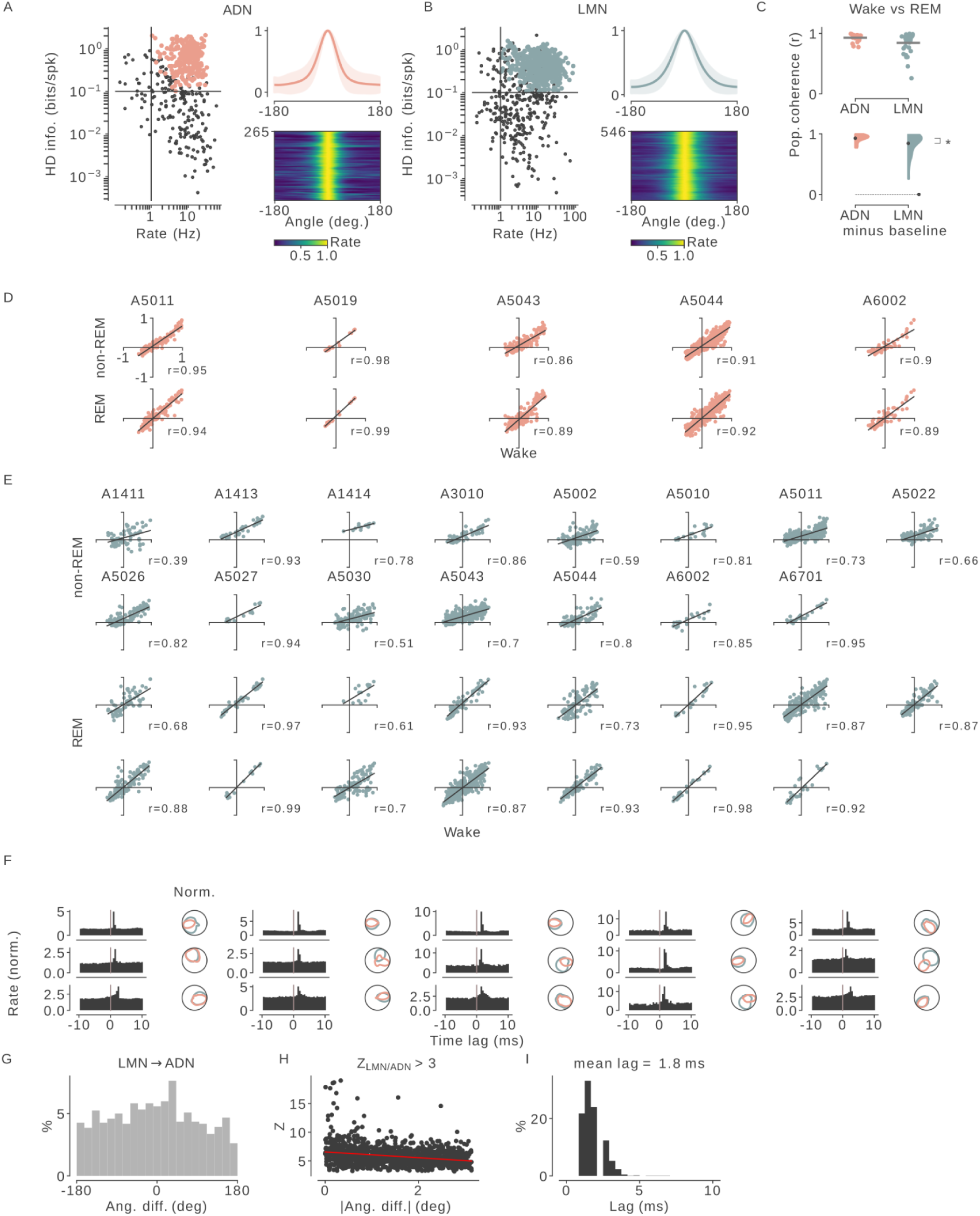
Population coherence of LMN and ADN units across individual sessions. **(A)** Left: Selection criteria for ADN units. Only cells with mean firing rates >1 Hz and HD information >0.1 bits/spike were included. Right: Tuning curves of all selected HD units in ADN. **(B)** Same as (A), for LMN units. **(C)** Population coherence during REM sleep for LMN and ADN units. Coherence was significantly higher in LMN (Mann–Whitney U = 276, p = 0.026; n_ADN_ = 24, n_LMN_ = 35). **(D)** Population coherence of ADN units across wakefulness and non-REM sleep, shown per session. **(E)** Same as (D), for LMN units. **(F)** Representative LMN→ADN cross-correlograms showing significant peaks (z > 3) within the 1–8 ms range. **(G)** Distribution of angular differences between preferred head directions of all significant LMN→ADN pairs. **(H)** Peak z-scores between 1 and 8 ms for all significantly interacting pairs, plotted as a function of the absolute angular difference of their preferred directions. **(I)** Distribution of time lags for all significantly interacting LMN→ADN pairs.

**Supplementary figure 3.**
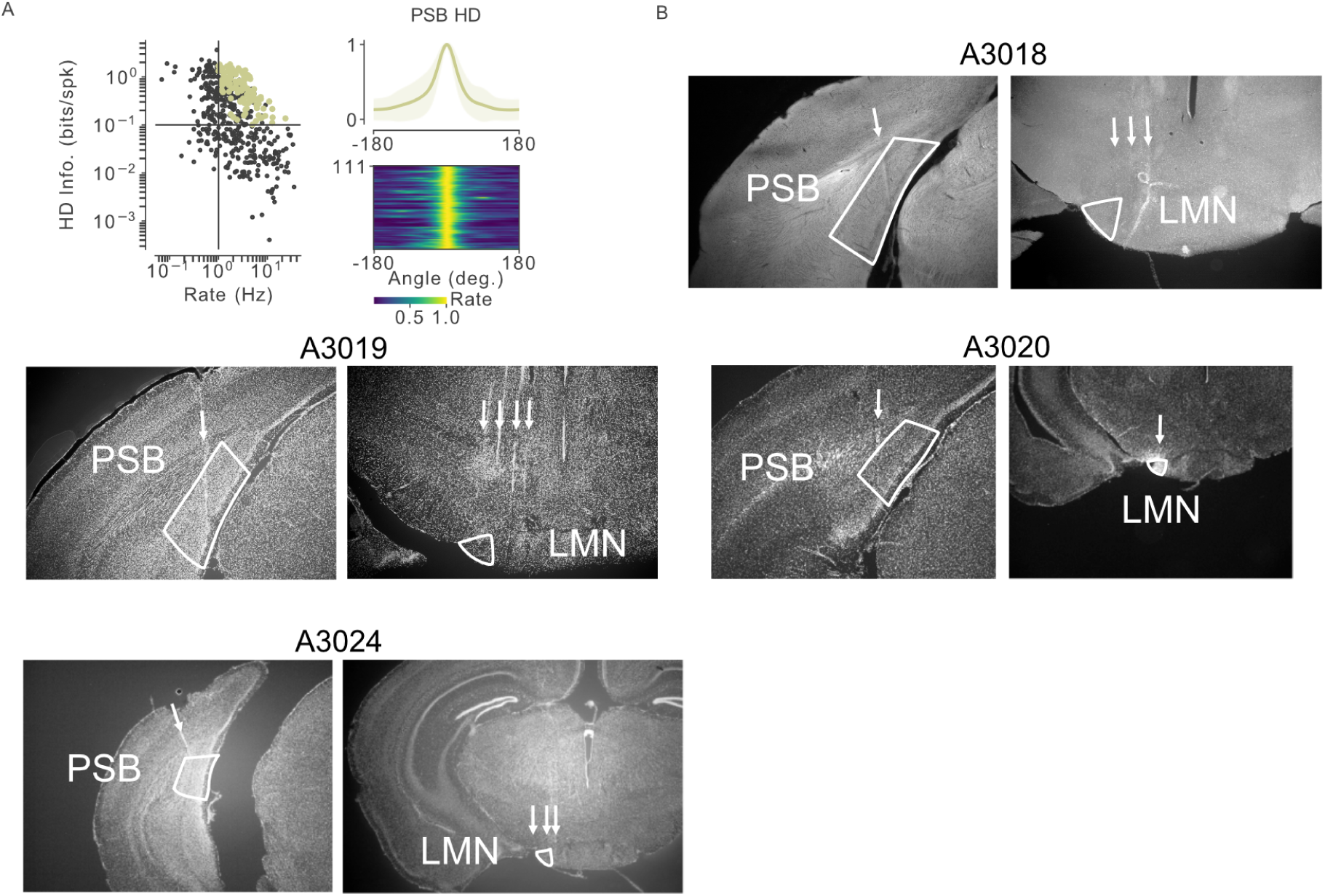
PSB and LMN double recordings. **(A)** Left: PSB units selection. Only units with rate larger than 1 Hz and HD information larger than 0.1 bits/spk were selected. Right: Tuning curves for all HD-PSD units selected. **(B)** Coronal histology showing silicon probe tracks (arrows) targeting the LMN and PSB.

**Supplementary Figure 4.**
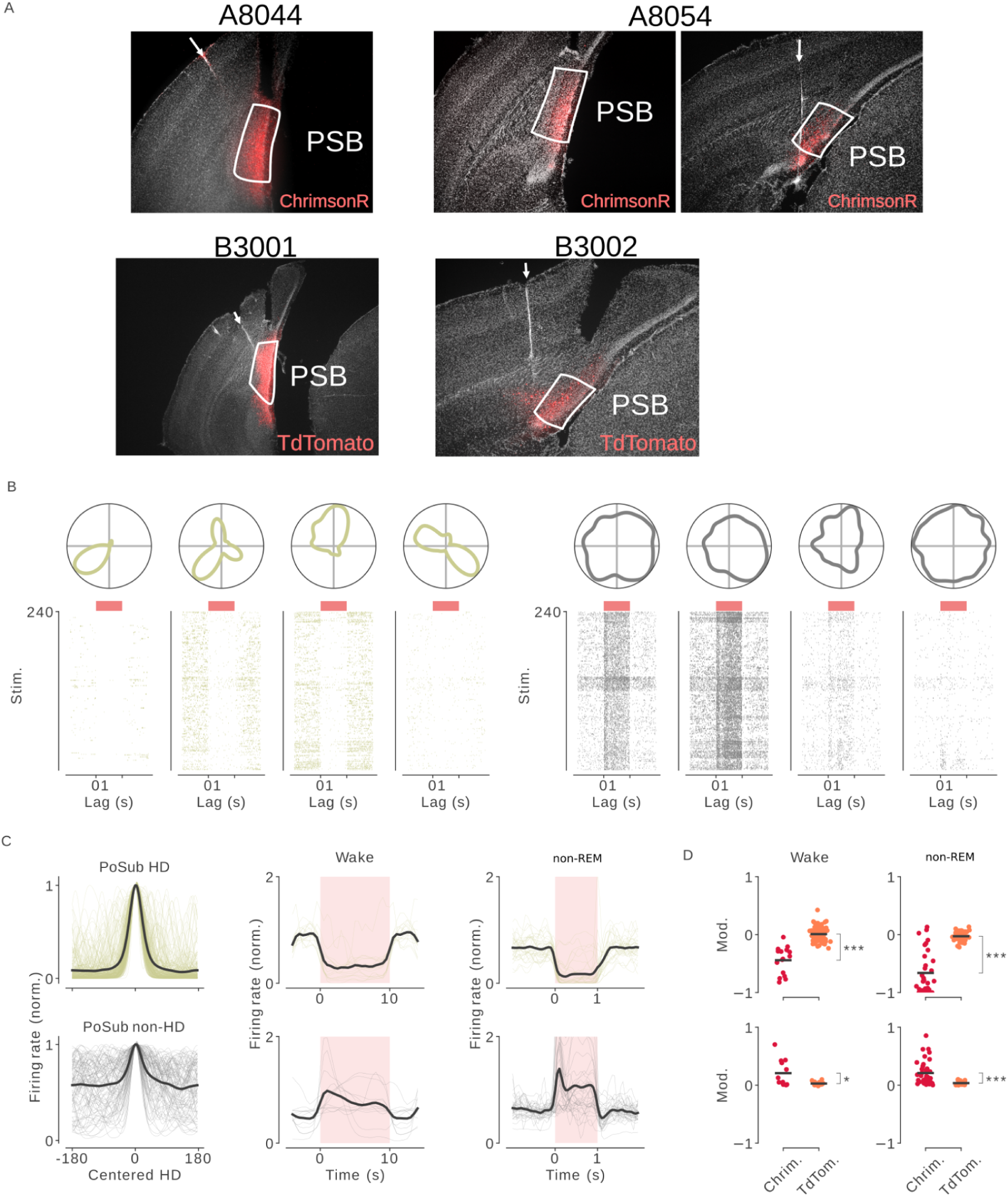
PSB injections and recordings in VGAT-Cre mice. **(A)** Coronal sections showing viral expression of ChrimsonR (n = 2 mice) or TdTomato (n = 2 mice) in the PSB, along with silicon probe placement. Arrows indicate probe tracks. **(B)** Example tuning curves and spike rasters for four HD cells and four non-HD cells recorded in the PSB during optogenetic stimulation in ChrimsonR-injected VGAT-Cre mice. **(C)** Mean-centered tuning curves for HD cells (top) and non-HD cells (bottom) during optogenetic stimulation. **(D)** Mean firing rates of HD and non-HD cells during wakefulness and non-REM sleep, under optogenetic stimulation. **(E)** Firing rate modulation in response to optogenetic stimulation, comparing ChrimsonR- and TdTomato-injected animals. For HD cells: Mann-Whitney; U_Wake_=36; p<0.001; U_Sleep_=266; p<0.001. For non-HD cells: Mann-Whitney; U_Wake_=195; p=0.02; U_Sleep_=813; p<0.001.

**Supplementary Figure 5.**
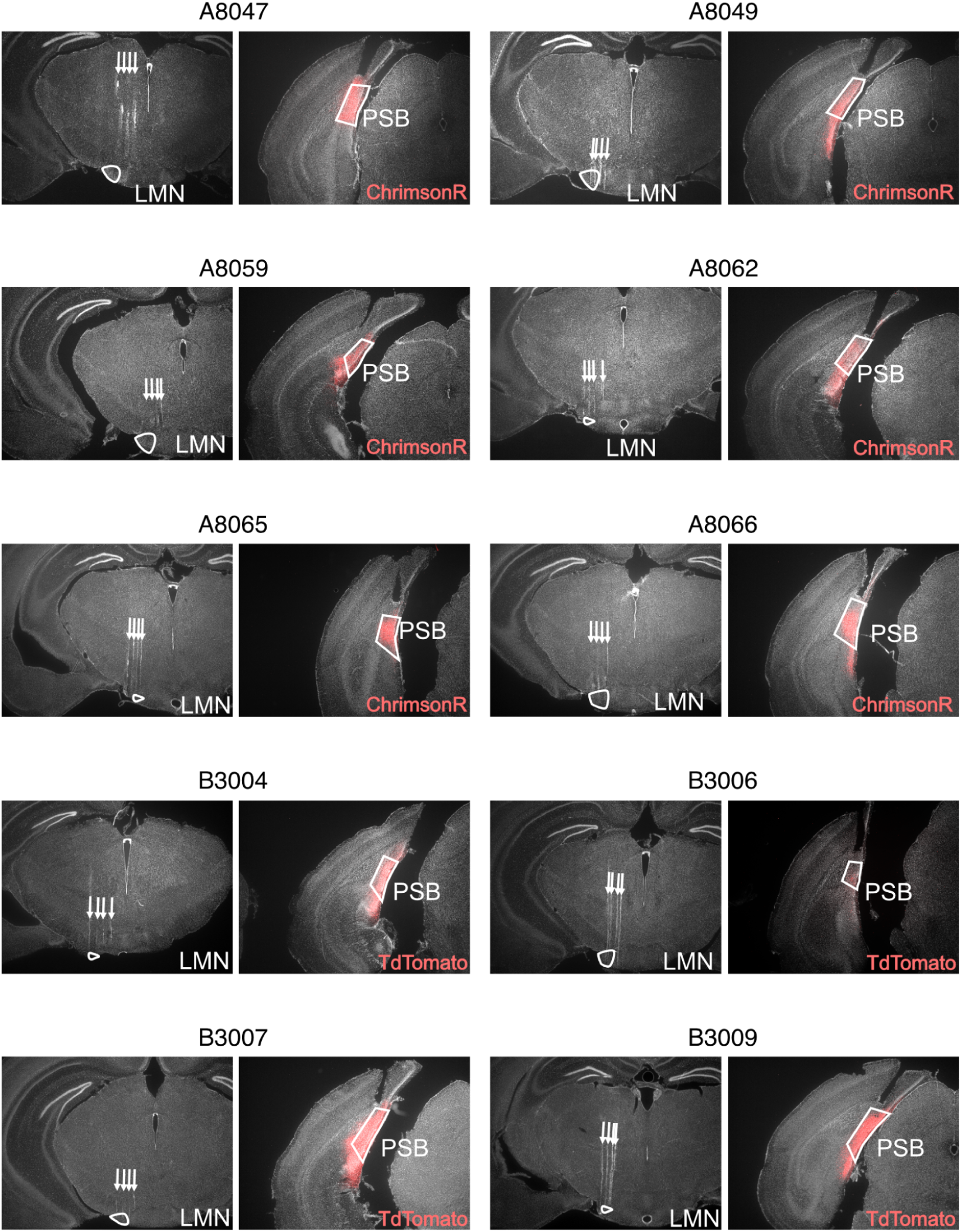
Histological verification of PSB injections and LMN recordings in VGAT-Cre mice. For each animal, coronal sections showing LMN (left) and PSB (right) histology are presented. Six mice were injected with ChrimsonR, and four control mice received TdTomato injections. Red labeling indicates the extent of viral expression.

**Supplementary Figure 6.**
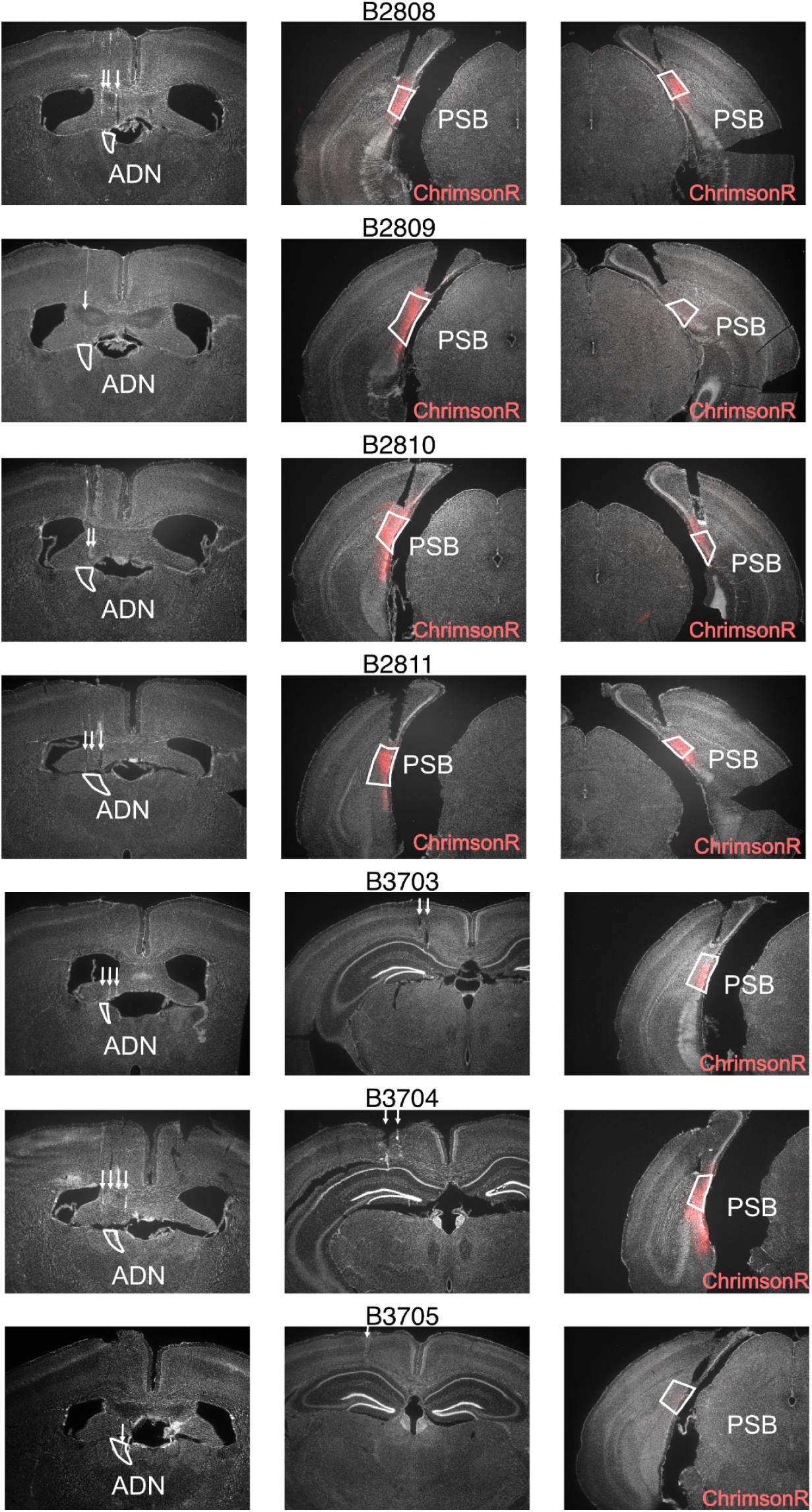
Histological verification of PSB injections and ADN recordings in VGAT-Cre mice. Red regions indicate viral expression. Mice B2809, B2810, and B2811 were injected bilaterally, while B3703, B3704, and B3705 received unilateral injections.

**Supplementary Figure 7.**
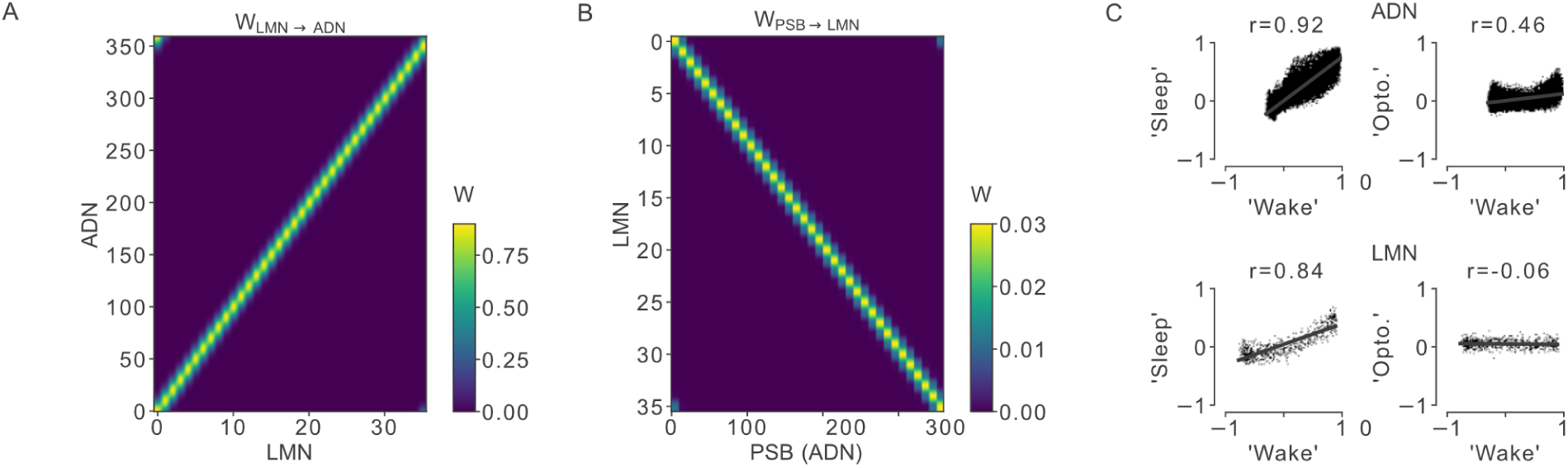
Network model details. **(A)** Connectivity matrix for LMN→ADN projections. **(B)** Connectivity matrix for PSB(ADN)→LMN projections. **(C)** Population coherence of LMN and ADN units across conditions: ‘wakefulness’ vs. ‘sleep’ and ‘wakefulness’ vs. ‘optogenetic stimulation’.

